# The impact of body posture on intrinsic brain activity: the role of beta power at rest

**DOI:** 10.1101/671818

**Authors:** Brunella Donno, Daniele Migliorati, Filippo Zappasodi, Mauro Gianni Perrucci, Marcello Costantini

**Author notes:** Corresponding authors: Donno Brunella., Laboratory of Neuropsychology and Cognitive Neuroscience, Department of Neuroscience, Imaging and Clinical Sciences, University G. d’Annunzio, Chieti, Italy & Institute for Advanced Biomedical Technologies (ITAB), University G. d’Annunzio, Chieti, Italy. Address: Via dei Vestini 31, 66100 Chieti (CH)., Or Costantini Marcello. Institute for Advanced Biomedical Technologies (ITAB), University G. d’Annunzio, Chieti, Italy. Address: Via dei Vestini 31, 66100 Chieti (CH).

## Abstract

Tying the hands behind the back has detrimental effect of sensorimotor perceptual tasks. Here we provide evidence that beta band oscillatory activity in a resting state condition might have a crucial role in such detrimental effects. EEG activity in a resting state condition was measured from thirty participants in two different body posture conditions. In one condition participants were required to keep their hands freely resting on the table. In the other condition, participants were required to keep the hands tied behind their back. Increased beta power was observed in the left inferior frontal gyrus (l-IFG) during the tied hands condition compared to the free hands condition. A control study ruled out alternative explanations including muscle tension that might have affected the EEG data. Our findings provide new insight on how body postural manipulations impact on perceptual tasks and intrinsic brain activity.

## 2. Introduction

It is well known that the physical body plays a key role in the way by which the brain encodes the environment; in fact, in everyday life, cognitive processes are influenced by the sensory and motor experiences of our body. This theory of cognition, known under the general topic of Embodied Cognition [1], claims that many aspects of cognition are shaped by features of the body [2]. This implies that if knowledge is obtained through bodily experiences, it is constrained not only by the experiences and situations encountered, but also by the physical features of the individuals [3]. A clear example is provided by studies on mental rotation of body parts [4-7]. In these tasks, participants judged the laterality of pictures representing hands and feet while standing in two different postural conditions. In one condition, the subjects’ right hand was placed on the right knee and the left hand behind the back; in the other one, the hand position was reversed. For right-handed subjects response times increased when participants judged images representing the right hands keeping their right hand behind the back. This was not found for images of the left hand and not for images of the feet. Other studies shown the same results highlighting an interference of hand posture with the ability to perform mental rotations of hand images [8]. Similarly, subjects’ body orientation has been shown to affect perception of both static and moving objects [9]. All these results suggest that information regarding the current positioning of body or body parts is required for an accurate encoding of visual information. Furthermore, it has been also demonstrated how the access to and retention of autobiographical memories in younger and older adults are affected by body posture [10]. In fact, response times were shorter when body positions during prompted retrieval of autobiographical events were similar to the body positions in the original events than when body position was incongruent [10]. Nevertheless, how the body posture impacts visual encoding of actions is still under debate. One possibility to understand how the body posture impacts visual encoding of actions is by looking at the interaction between posture manipulations and the intrinsic brain activity. Intrinsic brain activity is spontaneously generated by the brain and is not organized in a casual way [11]. The interaction between intrinsic brain activity and posture manipulations is better explained by a study of Thibault and collegues [12]. In this study prominent alterations of intrinsic brain activity over occipital and frontal regions have been shown due to orthostatic manipulations. Specifically, an increase of beta and gamma activity was observed while participants lied supine compared to the condition in which they either stand or sat inclined at 45 degrees. Moreover, interestingly strong pieces of evidence have shown that the intrinsic activity plays a key role in perceptual processes [13] involving high-frequency bands (i.e. beta and gamma) [14-16]. For instance, a decrease of oscillatory beta power has been observed over the sensorimotor regions during tasks requiring mental simulation of actions [17]. Such a decrease reflects the engagement of the motor system corresponding to the disinhibition of motor areas [18]. Indeed, an increased beta power has been shown to reflect an inhibition mechanisms related to perceptual and motor systems [19-22]. It is therefore conceivable that postural manipulations impact visual perception by altering beta-band oscillations. Drawing from this, we investigated the effects of postural manipulations on the intrinsic brain activity, focusing on the beta frequency band. EEG activity was measured in a resting state condition from thirty healthy participants in two body posture conditions. In one condition, participants were required to keep their hands freely resting on the table. In the other condition, participants were required to keep hands tied behind their back. Moreover, we conducted an EEG-EMG control experiment in order to rule out the presence of confounding variable (i.e. muscle tension). Specifically, subjects were asked to contract and to keep the contraction of specific muscles during the tied hands condition.

## 3. Materials and Method

### 3.1 Participants

Thirty right-handed healthy participants (12 males, mean age= 24.03; SD= 3.2; range= 20-33) were recruited to take part in the study from the student pool. The participants took part in the experiment at ITAB (Institute for Advanced Biomedical Technologies) in Chieti. Participants did not have any personal or close family history of neurological or psychiatric disorders, any brain surgery and any active medication, as self-reported. The study was approved by the local ethics committee (ID07022013). Participants gave their informed consent before taking part in the study. The study was conducted in accordance with the ethical standards of the 1964 Declaration of Helsinki.

## 3.2 Procedure

### 3.2.1 EEG resting-state recording and pre-processing

All participants underwent two different conditions (within-subject design): i) EEG resting-state when their hands were free (free condition) and ii) EEG resting-state when their hands were tied behind the back (tied condition). The two conditions were randomized across subjects. In the two blocks, participants had to keep their eyes open and fixate a cross in front of them placed on the computer screen. For each block, we recorded the neural activity in a resting state condition for 4 minutes.

We used a 64 electrodes cap (model BrainCap, BrainAmp MR Plus amplifier, Brain Products), placed according to the 10-20 International System. We used 2 electro-oculographic channels on the right and left temple to monitor eye movement and for off-line artefact rejection. The reference electrode was positioned in correspondence of FCz electrode while the ground electrode was placed in the Inion (Iz).

The impedance was measured before each recording and was kept below 10 kΩ. All the data were processed using EEGLAB software implemented in MATLAB [23].

We acquired online data at 5 kHz, band-pass filtered from 0.016 to 250 Hz. Data were off-line filtered between 1 Hz to 30 Hz and downsampled at 250 Hz for the current analysis. We detected and removed bad channels using a threshold with a probability at 5 % (*pop rejchan*). Subsequently, we rejected the continuous data using a threshold with a probability at 10 % (*pop rejcont*).

Finally, we computed the Independent Component Analysis, using the FastICA algorithm [24] to identify and reject manually noise, ocular, cardiac, muscular artefacts and bad channels. At this point, we interpolated rejected channel and EEG signal was re-referenced to the common average reference.

To exclude that differences in the beta band power in the tied hands with respect to free hands condition is not a consequence of muscular tension during the condition of tied hands, a control experiment was done. We co-registered EEG-EMG resting-state activity in 4 subjects (3 males, mean age= 26.75; SD=3.6; range= 24-32) during a low-level isometric contraction of the muscles for 4 minutes, recorded along with the EMG. During the tied hands condition, we asked to participants to contract deltoids, triceps, pectorals and dorsal muscles and to keep the contraction for 4 minutes as stable as possible.

Specifically, we recorded the contraction through 8 electromyographic channels: right and left deltoid, right and left pectoral, right and left dorsal, right and left triceps. A 32 electrodes cap (model BrainCap, BrainAmp MR Plus amplifier, Brain Products) was used. As the aim of the recording was to check the muscular artefact topography over the scalp, only ocular, cardiac and movement artifacts was rejected by ICA procedure. We computed the power spectrum density only for the beta band. The aim was to confirm a qualitative difference between the beta power spectrum scalp topography of the difference between tied versus free hands condition, obtained in the main experiment and the difference between the beta power scalp topography of the difference between contraction (tied hands condition) versus free hands condition, obtained in the control experiment. For the control experiment, for each subject we performed the beta power spectrum scalp topography of the difference between the two conditions.

## 3.3 EEG data analysis

### 3.3.1 Main experiment

We computed the power spectrum density for all electrodes using the periodogram Welch procedure (Hamming windowing function; window length 4 seconds; no overlap). The four classical EEG frequency bands were considered (delta: 1-4 Hz, theta: 4-8 Hz, alpha: 8-13 Hz and beta: 13-30 Hz). Delta, theta and alpha bands were used to control that the difference tied versus free hands condition was specific for the beta band.

Then we extracted the power of delta, theta, alpha and beta bands calculating the mean values of the power spectrum for all frequency bands and for the two conditions (free and tied) described above. The mean values were transformed into decibel scale (10 × log10[μV^2^]) in order to normalize the data [25].

To establish whether there were significant differences in power for all frequency bands between two conditions, a non-parametric cluster-based permutation test was performed using FieldTrip toolbox in MATLAB [26]. To investigate cortical generators of electrophysiological oscillations, we computed signal source localization for beta power spectral data, as this is the only frequency band showing differences between experimental conditions. The exact low resolution brain electromagnetic tomography (eLORETA) method in frequency domain was used to compute the cortical three-dimensional distribution of current density [27]. Computations are made in a realistic head model [28] using the Montreal Neurological Institute (MNI) Colin27 T2 template obtained from BrainWeb, (http://www.bic.mni.mcgill.ca/brainweb/).

Starting from estimated cortical distribution of generators of beta electrophysiological oscillations, the analysis of differences between free and tied hands condition showed a specific modulation whose significant neural source is localized in a specific cortical area.

## 3.4 EEG results

### 3.4.1 Main experiment

To test our hypothesis, we performed a non-parametric cluster-based permutation test for the power of all frequency bands between the two conditions (free and tied).

For each sample, a dependent-sample *t*-value was computed. All samples were selected for which this *t*-value exceeded an a priori threshold (uncorrected *p* < 0.05) and these were subsequently spatially clustered. The sum of the *t* values within a cluster was used as the cluster-level statistic. A reference distribution of cluster *t*-values was obtained by randomization of data across the two conditions for 5000 times and was used to evaluate the statistic of the actual data.

The non-parametric cluster-based permutation test revealed a significant difference between free and tied condition only in the beta band power (*ps* = 0.02). The analysis revealed increased beta power in tied hands condition compared to free hands condition and the difference between these two conditions was most pronounced over left inferior frontal electrodes (see Table 1 and Figure 1 for more details).

**Table 1:** Fronto-central electrodes in the significant cluster in beta power band (decibel: 10 × log10[μV^2^]) in the two experimental conditions (*ps* = 0.02).

| Electrodes | Free condition |  | Tied condition |  |
| --- | --- | --- | --- | --- |
|  | Mean | SD | Mean | SD |
| **F3** | -14,44 | 3,16 | -13,63 | 4,25 |
| **C3** | -15,89 | 2,94 | -15,04 | 3,96 |
| **F7** | -14,50 | 2,72 | -13,60 | 3,87 |
| **FC1** | -15,30 | 3,26 | -14,48 | 3,86 |
| **C1** | -16,72 | 3,07 | -15,95 | 3,75 |
| **FC3** | -15,58 | 3,13 | -14,68 | 3,68 |
| **F5** | -14,18 | 2,57 | -13,13 | 3,30 |
| **FT7** | -13,74 | 2,86 | -13,15 | 3,28 |

**Figure 1:**
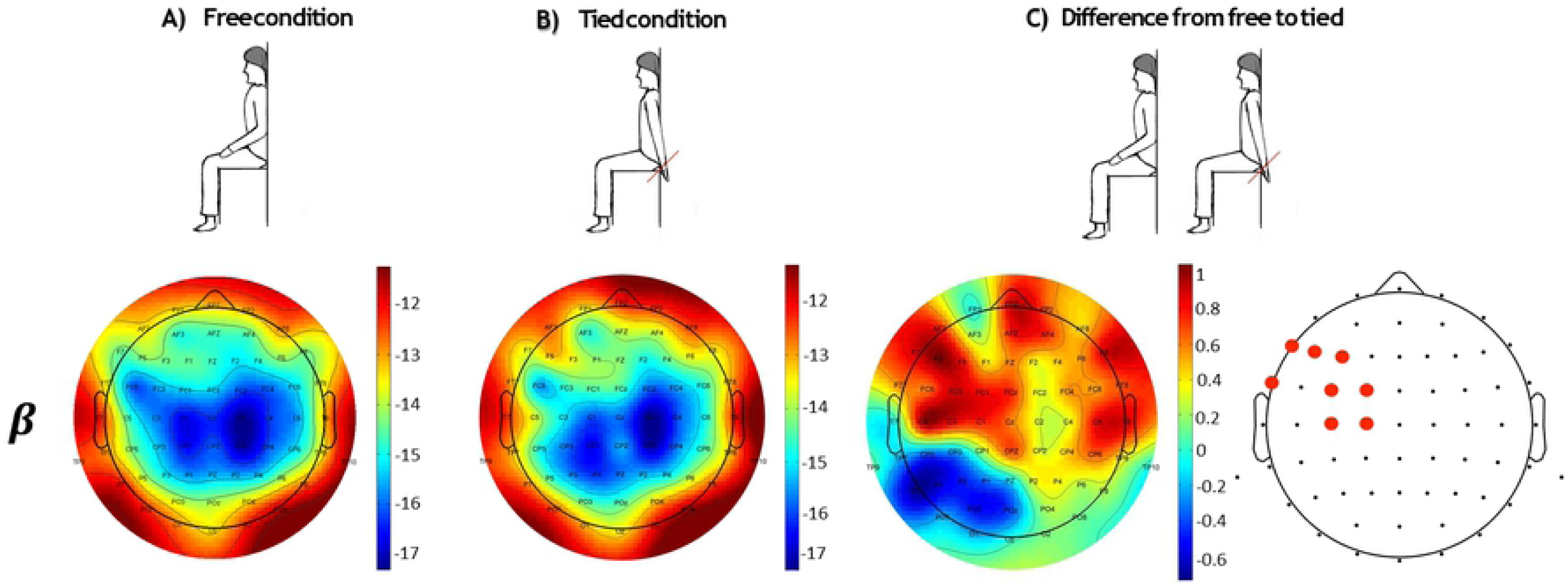
Scalp distributions of average beta power (decibel: 10 × log10[μV^2^]) in the free condition (Panel A) and tied condition (Panel B). Panel C represents the differences in beta power between the two conditions. Red dots represent electrodes in the significant cluster (*ps* = 0.02).

The non-parametric cluster-based permutation test did not reveal any significant differences between free and tied condition in the other frequency bands (all *ps* > 0.05). As regards signal source localization, the comparison between electrophysiological activity for beta power between free and tied hands conditions showed that the main signal source was in the left inferior frontal gyrus (l-IFG) (MNI: *x* = −35, *y* = 10, *z* = 15; *t* = 7.21) (See Figure 2).

**Figure 2:**
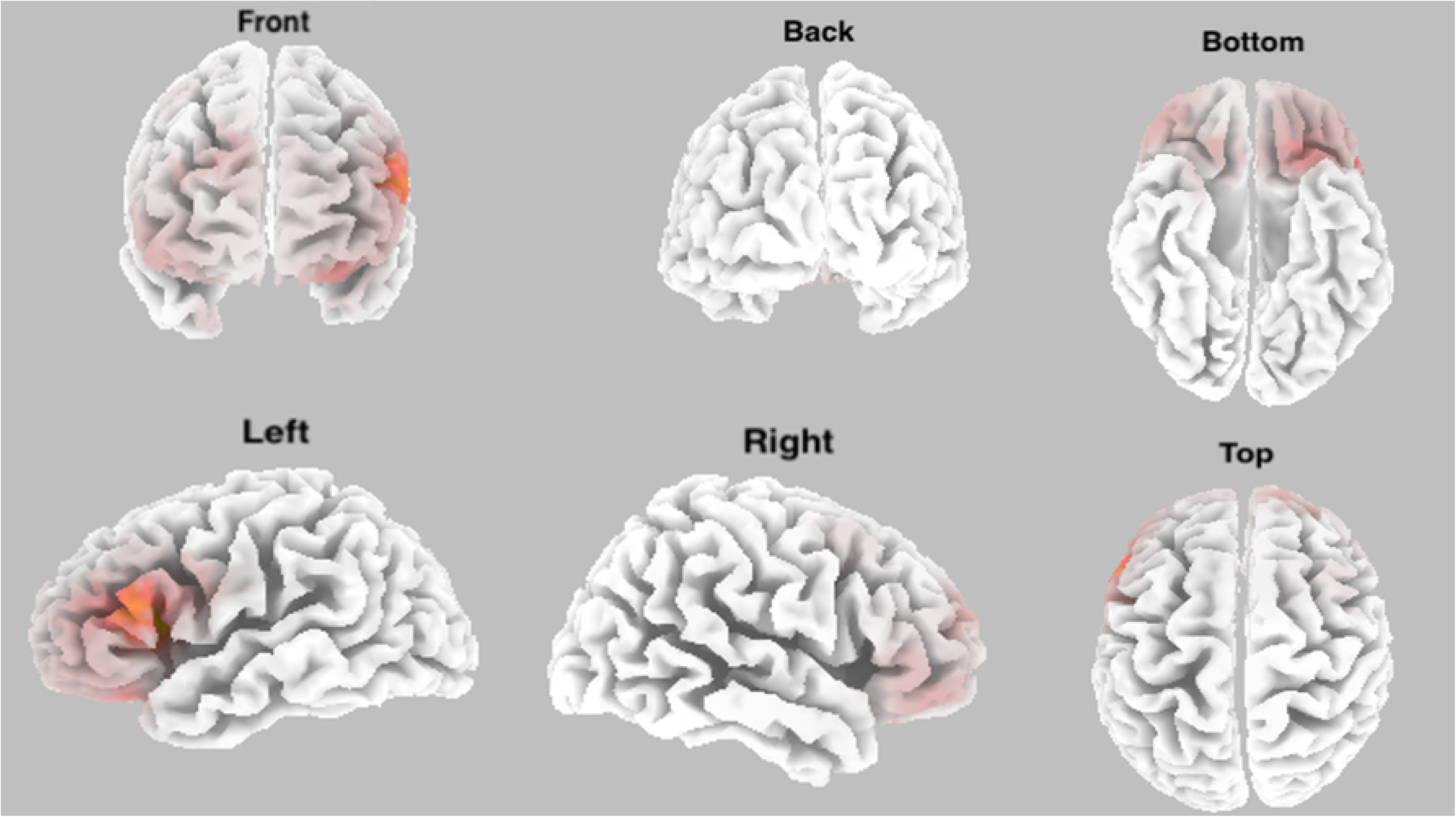
Source localization for beta power spectral data between free hands and tied hands conditions: eLORETA best match. Three-dimensional model reconstruction.

### 3.4.2 Control experiment

To exclude that differences in the beta band power in the tied hands with respect to free hands condition is not a consequence of muscular tension during the condition of tied hands, we compared the scalp distributions of average beta power relating to the difference between tied versus free hands condition, obtained in the main experiment and the scalp distributions of average beta power relating to the difference between contraction (tied hands condition) versus free hands condition, obtained in the control experiment. For each subject, we performed the beta power spectrum scalp topography of the difference between contraction (tied hands condition) versus free hands condition.

From each scalp topography, the maximum of the muscular artefact was located in the temporal, fronto-polar and parietal regions. In particular, muscular contamination was not present over the EEG channels where a significant difference between tied and free hands condition was found in the main experiment (F3, C3, F7, FC1, C1, FC3, F5, FT7) (See Figure 3).

**Figure 3:**
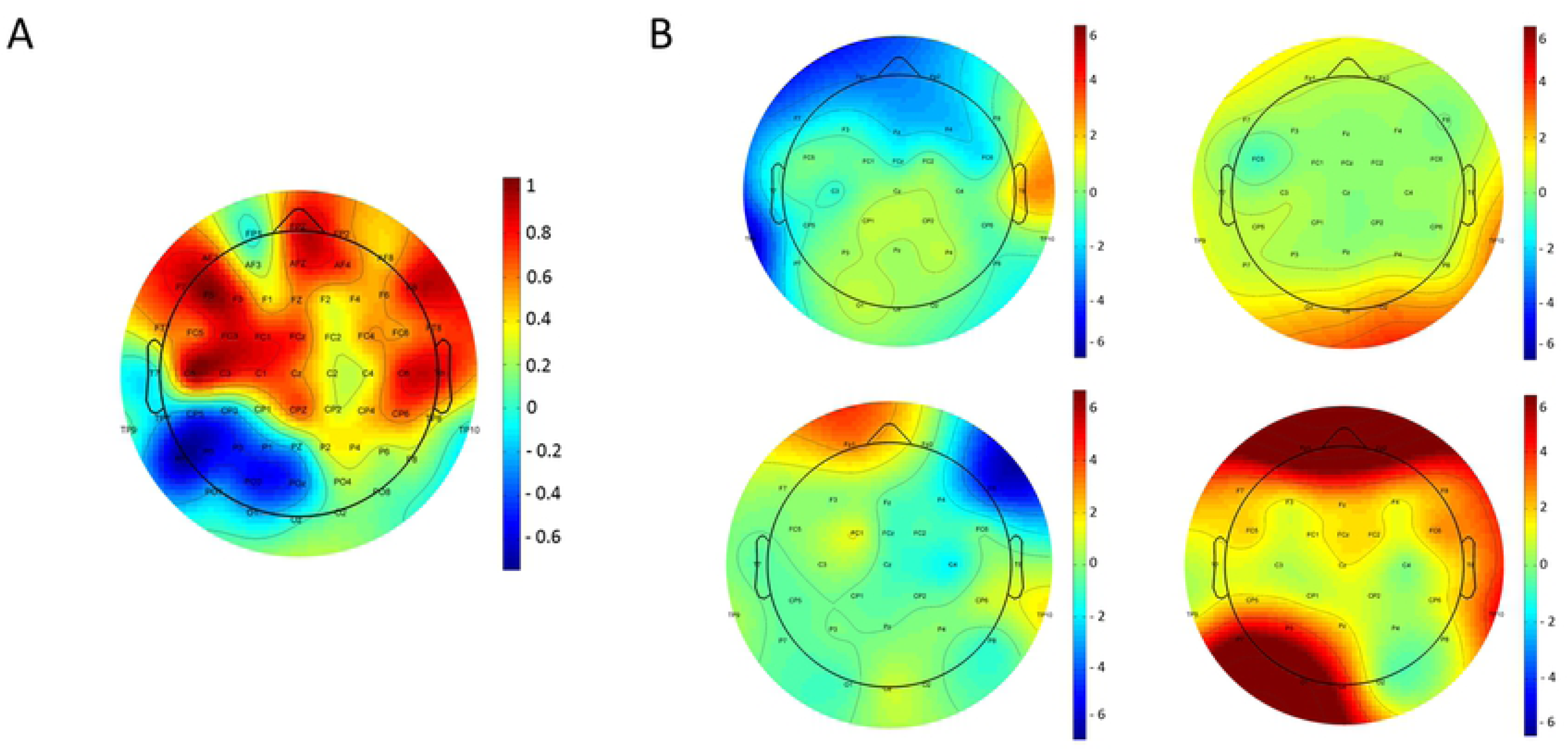
Scalp distributions of average beta power (decibel: 10 × log10[μV^2^]) of difference between tied versus free condition (Panel A) and 4 scalp distributions of average beta power (decibel: 10 × log10[μV^2^]) of difference between contraction (tied hands condition) versus free condition for each subject (Panel B).

Moreover, a similar topography was not found by visual inspection of beta band topographies of single subjects before the ICA algorithm application for artefact removal.

## 4. Discussion

We investigated the effect of body posture manipulations on the intrinsic brain activity. EEG activity in a resting state condition was measured from thirty healthy participants in two body posture conditions. In the free hands condition, participants were required to keep their hands freely resting on the table; in the tied hands condition, participants were required to keep hands tied behind their back. Power spectrum analysis revealed an increased beta power in the tied hands condition compared to the free hands condition. This difference was most pronounced over left frontal electrodes.

In this regard, to rule out the presence of confounding variables, such as increased muscle tension during tying hands at the back that could explain the observed beta band differences, we conducted an EEG-EMG control experiment. The two conditions were the same as the main experiment (free and tied hands). Specifically, during the tied hands condition we asked to participants to contract deltoids, triceps, pectorals and dorsal muscles and to keep the contraction as stable as possible. EEG activity in a resting state condition and EMG muscle activity in a resting state condition were measured from four participants.

It is well known what is the modulatory effect of body posture manipulations on sensorimotor perceptual tasks as well as it is known the involvement of beta power in the inhibition and disinhibition of motor mechanisms; nevertheless, what is the impact of the body posture on beta band oscillatory activity in a resting state condition is still unknown. Our results suggest that the increased beta power in the tied hands condition, compared to free hands condition, might explain how the constrained hands effect commonly observed when participants perform perceptual action-related tasks. Furthermore, regardless of the origin of the observed effect (muscle tension during the tying hands), we have shown how the neural outcome (the increase in beta power in the tied hands with respect to free hands condition) is not a consequence of muscular tension during the condition of tied hands. In this regard, we compared the scalp topography of beta power distribution of the control experiment and the main experiment. To confirm that the data obtained was not a consequence of muscular tension, we compared the scalp topography of beta power distribution relating to the difference between contraction (tied hands condition) versus free hands condition for each subject obtained during the control experiment and the topography of the average beta power distribution relating to the difference between tied versus free hands condition obtained during the main experiment. Regarding the data obtained, our results might explain clearly how the constrained hands effect commonly observed when participants perform perceptual action-related tasks. But really does beta power activity in sensorimotor regions play a role in processing such stimuli? More, how strong its involvement is in their processing? Previous studies have shown that observation of graspable objects, which is known to recruit sensorimotor resources [29] and is known to be affected by postural manipulations [21], is associated with a decrease in beta band power. Similarly, suppression of oscillatory activity within the mu (8–13 Hz) and beta (13–30 Hz) frequency bands over sensorimotor regions has been associated with action execution, as well as action observation [30-34]. Moreover, it has been similarly found how also passive observation of manipulable objects elicits similar neural responses to passive observation of others’ actions [29, 35-38], as well as in an EEG study, that examined neural activity during processing of tools, showed such suppression of beta activity [39]. Hence, increasing evidence suggests that the power of beta rhythm is typically decreased during the preparation and execution of a movement [40]; it increases in the motor cortex during active immobilization [41], postural maintenance [42], proactive inhibition [43], when a movement have to be withheld or voluntary suppressed [44] but also before an expected postural challenge [45]. Moreover, interestingly strong pieces of evidence have demonstrated that beta power enhancement with transcranial alternating cortical stimulation has been shown to induce motor inhibition [46].

Similar results have been found when rhythmic activity is induced in the motor cortex of healthy participants using transcranial current stimulation, where the stimulation in the beta band frequency range, that reflects an increased beta power, is particularly effective in slowing movements and increasing the threshold of inducing a motor response [47-49].

But the role played by the beta band in inhibition/disinhibition of neural motor system is supported also by studies on clinical populations. Specifically, the functional relevance of the beta band rhythm in the disinhibition of neuronal populations becomes particularly clear in Parkinson’s disease (PD), where pathological high beta band activity severely compromises movement initiation and execution [50, 51]. More generally, these findings support the idea that the beta band power maintains the functioning of the sensorimotor cortex [47, 48]. Taken together, these data are compatible with the hypothesis that beta band activity may signal the tendency of the sensorimotor system to maintain the status quo [52]. An interesting suggestion in this context is that beta band activity may allow the more efficient processing of feedback (e.g. proprioceptive signals) that is required for monitoring the status quo and recalibrating the sensorimotor system [52].

Furthermore, to investigate cortical generators of electrophysiological oscillations of beta frequency band, we performed signal source localization for beta power spectral data. The comparison between electrophysiological activity in the free hands and tied hands condition showed that the main signal source was localised in the left inferior frontal gyrus (l-IFG). The involvement of the l-IFG in processing action related stimuli has been shown by a large amount of studies. For instance, it has been shown that this cortex is critically involved not only in planning and executing object-related hand actions [53, 54], but also in processing both others’ object-related actions and action-related features of objects. Moreover, a large number of studies have demonstrated that viewing another’s object related action recruits the left ventral premotor cortex (PMv) as if the viewer were performing that action herself [55-60]. Finally, this area has been shown to be involved in response inhibition in a Go/NoGo task, demonstrating how the integrity of this area is critical for successful implementation of inhibitory control over motor responses [61], as well as it is crucially involved in processing visual features of objects in terms of the actions they might afford [62-65]. In this context, it has been demonstrated how l-IFG and PMv are significantly activated during gesture planning and tool use actions [66]. To sum up, we have shown the effect of tying the hands on intrinsic brain activity and how this manipulation can change the activity in the beta frequency band in a resting state condition. Our result might contribute to explain the constrained hand effect commonly observed when participants perform perceptual action-related tasks.

## References

1. Shapiro L. Embodied cognition: Routledge; 2010.

2. Rowlands M. The new science of the mind: From extended mind to embodied phenomenology: Mit Press; 2010.

3. Wilson RA, Foglia L. Embodied cognition. 2011.

4. Bonda E, Petrides M, Frey S, EvANs A. Neural correlates of mental transformations of the body-in-space. Proceedings of the National Academy of Sciences. 1995;92(24):11180–4.

5. Cohen RG, Rosenbaum DA. Prospective and retrospective effects in human motor control: planning grasps for object rotation and translation. Psychological Research. 2011;75(4):341–9.

6. Ionta S, Perruchoud D, Draganski B, Blanke O. Body context and posture affect mental imagery of hands. PloS one. 2012;7(3):e34382.

7. Overney LS, Michel CM, Harris IM, Pegna AJ. Cerebral processes in mental transformations of body parts: recognition prior to rotation. Cognitive brain research. 2005;25(3):722–34.

8. Ionta S, Blanke O. Differential influence of hands posture on mental rotation of hands and feet in left and right handers. Experimental brain research. 2009;195(2):207–17.

9. Lopez C, Bachofner C, Mercier M, Blanke O. Gravity and observer’s body orientation influence the visual perception of human body postures. Journal of vision. 2009;9(5):1-.

10. Dijkstra K, Kaschak MP, Zwaan RA. Body posture facilitates retrieval of autobiographical memories. Cognition. 2007;102(1):139–49.

11. Fox MD, Snyder AZ, Vincent JL, Corbetta M, Van Essen DC, Raichle ME. The human brain is intrinsically organized into dynamic, anticorrelated functional networks. Proceedings of the National Academy of Sciences. 2005;102(27):9673–8.

12. Thibault RT, Lifshitz M, Raz A. Body position alters human resting-state: Insights from multi-postural magnetoencephalography. Brain imaging and behavior. 2016;10(3):772–80.

13. Pfurtscheller G, Aranibar A. Evaluation of event-related desynchronization (ERD) preceding and following voluntary self-paced movement. Electroencephalography and clinical neurophysiology. 1979;46(2):138–46.

14. Herrmann CS, Munk MH, Engel AK. Cognitive functions of gamma-band activity: memory match and utilization. Trends in cognitive sciences. 2004;8(8):347–55.

15. Martinovic J, Busch NA. High frequency oscillations as a correlate of visual perception. International Journal of Psychophysiology. 2011;79(1):32–8.

16. Tallon-Baudry C. The roles of gamma-band oscillatory synchrony in human visual cognition. Front Biosci. 2009;14:321–32.

17. Brinkman L, Stolk A, Dijkerman HC, de Lange FP, Toni I. Distinct roles for alpha-and beta-band oscillations during mental simulation of goal-directed actions. Journal of Neuroscience. 2014;34(44):14783–92.

18. Cheyne DO. MEG studies of sensorimotor rhythms: a review. Experimental neurology. 2013;245:27–39.

19. Hari R, Salmelin R. Human cortical oscillations: a neuromagnetic view through the skull. Trends in neurosciences. 1997;20(1):44–9.

20. Jensen O, Goel P, Kopell N, Pohja M, Hari R, Ermentrout B. On the human sensorimotor-cortex beta rhythm: sources and modeling. Neuroimage. 2005;26(2):347–55.

21. Natraj N, Poole V, Mizelle J, Flumini A, Borghi AM, Wheaton LA. Context and hand posture modulate the neural dynamics of tool–object perception. Neuropsychologia. 2013;51(3):506–19.

22. Zimmermann M, Toni I, de Lange FP. Body posture modulates action perception. Journal of Neuroscience. 2013;33(14):5930–8.

23. Delorme A, Makeig S. EEGLAB: an open source toolbox for analysis of single-trial EEG dynamics including independent component analysis. Journal of neuroscience methods. 2004;134(1):9–21.

24. Hyvärinen A, Oja E. Independent component analysis: algorithms and applications. Neural networks. 2000;13(4-5):411–30.

25. Fell J, Widman G, Rehberg B, Elger CE, Fernandez G. Human mediotemporal EEG characteristics during propofol anesthesia. Biological cybernetics. 2005;92(2):92–100.

26. Oostenveld R, Fries P, Maris E, Schoffelen J-M. FieldTrip: open source software for advanced analysis of MEG, EEG, and invasive electrophysiological data. Computational intelligence and neuroscience. 2011;2011:1.

27. Pascual-Marqui RD. Discrete, 3D distributed, linear imaging methods of electric neuronal activity. Part 1: exact, zero error localization. arXiv preprint arXiv:07103341. 2007.

28. Fuchs M, Kastner J, Wagner M, Hawes S, Ebersole JS. A standardized boundary element method volume conductor model. Clinical Neurophysiology. 2002;113(5):702–12.

29. Creem-Regehr SH, Lee JN. Neural representations of graspable objects: are tools special? Cognitive Brain Research. 2005;22(3):457–69.

30. Arnstein D, Cui F, Keysers C, Maurits NM, Gazzola V. μ-suppression during action observation and execution correlates with BOLD in dorsal premotor, inferior parietal, and SI cortices. Journal of Neuroscience. 2011;31(40):14243–9.

31. Cochin S, Barthelemy C, Lejeune B, Roux S, Martineau J. Perception of motion and qEEG activity in human adults. Electroencephalography and clinical neurophysiology. 1998;107(4):287–95.

32. Frenkel-Toledo S, Bentin S, Perry A, Liebermann DG, Soroker N. Dynamics of the EEG power in the frequency and spatial domains during observation and execution of manual movements. Brain research. 2013;1509:43–57.

33. Hari R, Forss N, Avikainen S, Kirveskari E, Salenius S, Rizzolatti G. Activation of human primary motor cortex during action observation: a neuromagnetic study. Proceedings of the National Academy of Sciences. 1998;95(25):15061–5.

34. Perry A, Bentin S. Mirror activity in the human brain while observing hand movements: A comparison between EEG desynchronization in the μ-range and previous fMRI results. Brain research. 2009;1282:126–32.

35. Caggiano V, Fogassi L, Rizzolatti G, Thier P, Casile A. Mirror neurons differentially encode the peripersonal and extrapersonal space of monkeys. science. 2009;324(5925):403–6.

36. Cisek P, Kalaska JF. Neural mechanisms for interacting with a world full of action choices. Annual review of neuroscience. 2010;33:269–98.

37. Proverbio AM. Tool perception suppresses 10–12Hz μ rhythm of EEG over the somatosensory area. Biological psychology. 2012;91(1):1–7.

38. Proverbio AM, Adorni R, D’Aniello GE. 250ms to code for action affordance during observation of manipulable objects. Neuropsychologia. 2011;49(9):2711–7.

39. Simon S, Mukamel R. Power modulation of electroencephalogram mu and beta frequency depends on perceived level of observed actions. Brain and behavior. 2016;6(8).

40. Pfurtscheller G, Da Silva FL. Event-related EEG/MEG synchronization and desynchronization: basic principles. Clinical neurophysiology. 1999;110(11):1842–57.

41. Salmelin R, Hámáaláinen M, Kajola M, Hari R. Functional segregation of movement-related rhythmic activity in the human brain. Neuroimage. 1995;2(4):237–43.

42. Baker S, Olivier E, Lemon R. Coherent oscillations in monkey motor cortex and hand muscle EMG show task-dependent modulation. The Journal of physiology. 1997;501(1):225–41.

43. Zavala B, Zaghloul K, Brown P. The subthalamic nucleus, oscillations, and conflict. Movement Disorders. 2015;30(3):328–38.

44. Swann N, Tandon N, Canolty R, Ellmore TM, McEvoy LK, Dreyer S, et al. Intracranial EEG reveals a time-and frequency-specific role for the right inferior frontal gyrus and primary motor cortex in stopping initiated responses. Journal of Neuroscience. 2009;29(40):12675–85.

45. Androulidakis AG, Doyle LM, Yarrow K, Litvak V, Gilbertson TP, Brown P. Anticipatory changes in beta synchrony in the human corticospinal system and associated improvements in task performance. European Journal of Neuroscience. 2007;25(12):3758–65.

46. Joundi RA, Jenkinson N, Brittain J-S, Aziz TZ, Brown P. Driving oscillatory activity in the human cortex enhances motor performance. Current Biology. 2012;22(5):403–7.

47. Feurra M, Bianco G, Santarnecchi E, Del Testa M, Rossi A, Rossi S. Frequency-dependent tuning of the human motor system induced by transcranial oscillatory potentials. Journal of Neuroscience. 2011;31(34):12165–70.

48. Pogosyan A, Gaynor LD, Eusebio A, Brown P. Boosting cortical activity at beta-band frequencies slows movement in humans. Current Biology. 2009;19(19):1637–41.

49. Wach C, Krause V, Moliadze V, Paulus W, Schnitzler A, Pollok B. Effects of 10Hz and 20Hz transcranial alternating current stimulation (tACS) on motor functions and motor cortical excitability. Behavioural brain research. 2013;241:1–6.

50. Brown P. Abnormal oscillatory synchronisation in the motor system leads to impaired movement. Current opinion in neurobiology. 2007;17(6):656–64.

51. Jenkinson N, Brown P. New insights into the relationship between dopamine, beta oscillations and motor function. Trends in neurosciences. 2011;34(12):611–8.

52. Engel AK, Fries P. Beta-band oscillations—signalling the status quo? Current opinion in neurobiology. 2010;20(2):156–65.

53. Jeannerod M, Arbib MA, Rizzolatti G, Sakata H. Grasping objects: the cortical mechanisms of visuomotor transformation. Trends in neurosciences. 1995;18(7):314–20.

54. Rizzolatti G, Gentilucci M. Motor and visual-motor functions of the premotor cortex. Neurobiology of neocortex. 1988;42:269–84.

55. Buccino G, Binkofski F, Fink GR, Fadiga L, Fogassi L, Gallese V, et al. Action observation activates premotor and parietal areas in a somatotopic manner: an fMRI study. European journal of neuroscience. 2001;13(2):400–4.

56. Calvo-Merino B, Grèzes J, Glaser DE, Passingham RE, Haggard P. Seeing or doing? Influence of visual and motor familiarity in action observation. Current Biology. 2006;16(19):1905–10.

57. Galati G, Committeri G, Spitoni G, Aprile T, Di Russo F, Pitzalis S, et al. A selective representation of the meaning of actions in the auditory mirror system. Neuroimage. 2008;40(3):1274–86.

58. Grafton ST, Arbib MA, Fadiga L, Rizzolatti G. Localization of grasp representations in humans by positron emission tomography. Experimental brain research. 1996;112(1):103–11.

59. Ortigue S, Sinigaglia C, Rizzolatti G, Grafton ST. Understanding actions of others: the electrodynamics of the left and right hemispheres. A high-density EEG neuroimaging study. PloS one. 2010;5(8):e12160.

60. Rizzolatti G, Fadiga L, Gallese V, Fogassi L. Premotor cortex and the recognition of motor actions. Cognitive brain research. 1996;3(2):131–41.

61. Swick D, Ashley V, Turken U. Left inferior frontal gyrus is critical for response inhibition. BMC neuroscience. 2008;9(1):102.

62. Buccino G, Sato M, Cattaneo L, Rodà F, Riggio L. Broken affordances, broken objects: a TMS study. Neuropsychologia. 2009;47(14):3074–8.

63. Chao LL, Martin A. Representation of manipulable man-made objects in the dorsal stream. Neuroimage. 2000;12(4):478–84.

64. Grafton ST, Fadiga L, Arbib MA, Rizzolatti G. Premotor cortex activation during observation and naming of familiar tools. Neuroimage. 1997;6(4):231–6.

65. Grèzes J, Tucker M, Armony J, Ellis R, Passingham RE. Objects automatically potentiate action: an fMRI study of implicit processing. European Journal of Neuroscience. 2003;17(12):2735–40.

66. Johnson-Frey SH, Newman-Norlund R, Grafton ST. A distributed left hemisphere network active during planning of everyday tool use skills. Cerebral cortex. 2004;15(6):681–95.

